# Novel fungal metal-dependent GH54 α-L-arabinofuranosidase: expanded substrate specificity and potential use for plant biomass degradation

**DOI:** 10.1101/2020.12.18.423520

**Authors:** Maria Lorenza Leal Motta, Jaire Alves Ferreira Filho, Ricardo Rodrigues de Melo, Leticia Maria Zanphorlin Murakami, Clelton Aparecido dos Santos, Anete Pereira Souza

## Abstract

*Trichoderma* genus fungi present great potential for the production of carbohydrate-active enzymes (CAZYmes), including glycoside hydrolase (GH) family members. From a renewability perspective, CAZYmes can be biotechnologically exploited to convert plant biomass into free sugars for the production of advanced biofuels and other high-value chemicals. GH54 is an attractive enzyme family for biotechnological applications because many GH54 enzymes are bifunctional. Thus, GH54 enzymes are interesting targets in the search for new enzymes for use in industrial processes such as plant biomass conversion. Herein, a novel metal-dependent GH54 arabinofuranosidase (ThABF) from the cellulolytic fungus *Trichoderma harzianum* was identified and biochemically characterized. Initial *in silico* searches were performed to identify the GH54 sequence. Next, the gene was cloned and heterologously overexpressed in *Escherichia coli*. The recombinant protein was purified, and the enzyme’s biochemical and biophysical properties were assessed. The GH54 members show wide functional diversity and specifically remove plant cell decorations including arabinose and galactose, in the presence of a metallic cofactor. Plant cell wall decoration have a major impact on lignocellulosic substrate conversion into high-value chemicals. These results expand the known functional diversity within the GH54 family, showing the potential of a novel arabinofuranosidase for plant biomass degradation.

## Introduction

Arabinofuranosidases (ABFs) (EC 3.2.1.55) are enzymes that are capable of cleaving residues of l-arabinofuranosyl present in various oligosaccharides and in polysaccharides such as hemicellulose; for this reason, they are interesting targets for biotechnological applications^1^. These enzymes are part of a group of glycosidases necessary for the degradation of polymeric substrates, such as arabinane, arabinoxylan and other polysaccharides, that constitute the walls of plant cells^2^. ABFs are grouped in glycosidic hydrolase (GH) families 3, 43, 51, 54 and 62, where GH54 includes some enzymes that are described as bifunctional^3^. However, few studies focusing on the functional specificity of this family have been conducted.

Ravanal *et al*.^4^ characterized an α-L-arabinofuranosidase from *Penicillium purpurogenum* and demonstrated that this family includes enzymes that are capable of acting synergistically with others, both releasing arabinose from different sources and acting on other substrates. Thus, given the features presented by this group of enzymes, further studies are still needed to better understand and biotechnologically explore the enzymes in this family, especially with regard to their application for the conversion of lignocellulosic substrates into bioproducts, such as biofuels.

Vegetable biomass is a reservoir of sugars organized in complex regular structures represented by cellulose, hemicellulose and lignin^5^. Such structures need to be deconstructed for the release of sugar-free monomers, which can then be converted into high-added-value chemicals such as second-generation ethanol^6^. This process requires the enzymatic conversion of biomass, and hemicellulose is a biopolymer that requires a large complex of enzymes for its conversion due to its structural composition^7^.

For the enzymes to access the main chain of hemicellulose, it is necessary to remove side-chain decorations, as they are one of the main reasons for the recalcintrance of plant biomass^8,9^. Thus, the search for enzymes that target hemicellulose decorations is a promising approach for improving the saccharification of lignocellulolytic material^10^. It is of interest to apply ABFs for this purpose because in addition, removing l-arabinose, some of these enzymes can act on other sugars involved in decorations^1^.

These enzymes are particularly important in the genus *Trichoderma*, where they play a crucial physiological role^11^. This genus is composed of heterotrophic filamentous ascomycetous fungi that are capable of growing on different substrates and under different environmental conditions. There are several studies that have addressed the versatility of this genus, which range from applications in sugar and alcohol industries to the development of biofungicides^12^. *Trichoderma reesei* is of particular prominence within the group because of its recognized potential to produce several hydrolytic enzymes; however, other species of the genus, especially *Trichoderma harzianum*, have been shown to be as efficient as *T. reesei* in this regard^13^.

Therefore, in this study, we conducted the *in silico* bioprospecting of a new GH54 from *T. harzianum*. For this purpose, several approaches were used, such as RNA-seq, in addition to sequence analysis. The enzyme was also characterized biochemically with different substrates and metallic salts to determine their influence on the enzyme.

## Results

### *Obtaining the sequence of ThABF and its* in silico *characterization*

Through database searches, a sequence from *Trichoderma koningii* (AAA81024.1) was selected and used in the subsequent steps detailed in the methodology. With the resulting BLAST sequence, the *T. harzianum CBS 226.95* genome was mapped the *T. harzianum* IOC-3844 RNA-seq reads and obtained the target protein sequence (MT439956). With this target protein sequence, a BLAST search was performed against the CAZy database, where our data suggest that ThABF is more similar to the GH54 family and that this family has several carbohydrate-binding models (CBMs; 1, 2, 6, 13 and 42), as shown in Supplementary Table S1.

A phylogenetic analysis of ThABF against sequences of the GH54 family (represented mainly by fungal and bacterial sequences) revealed great diversity between the sequences (Fig. 1a). The sequences used in the analysis are in Supplementary Table S2. The phylogenetic tree was divided into five groups that best explained this diversity: a Fungi group, containing only fungi enzymes, and Bacterium I to IV groups, containing bacterial enzymes.

**Figure 1.**
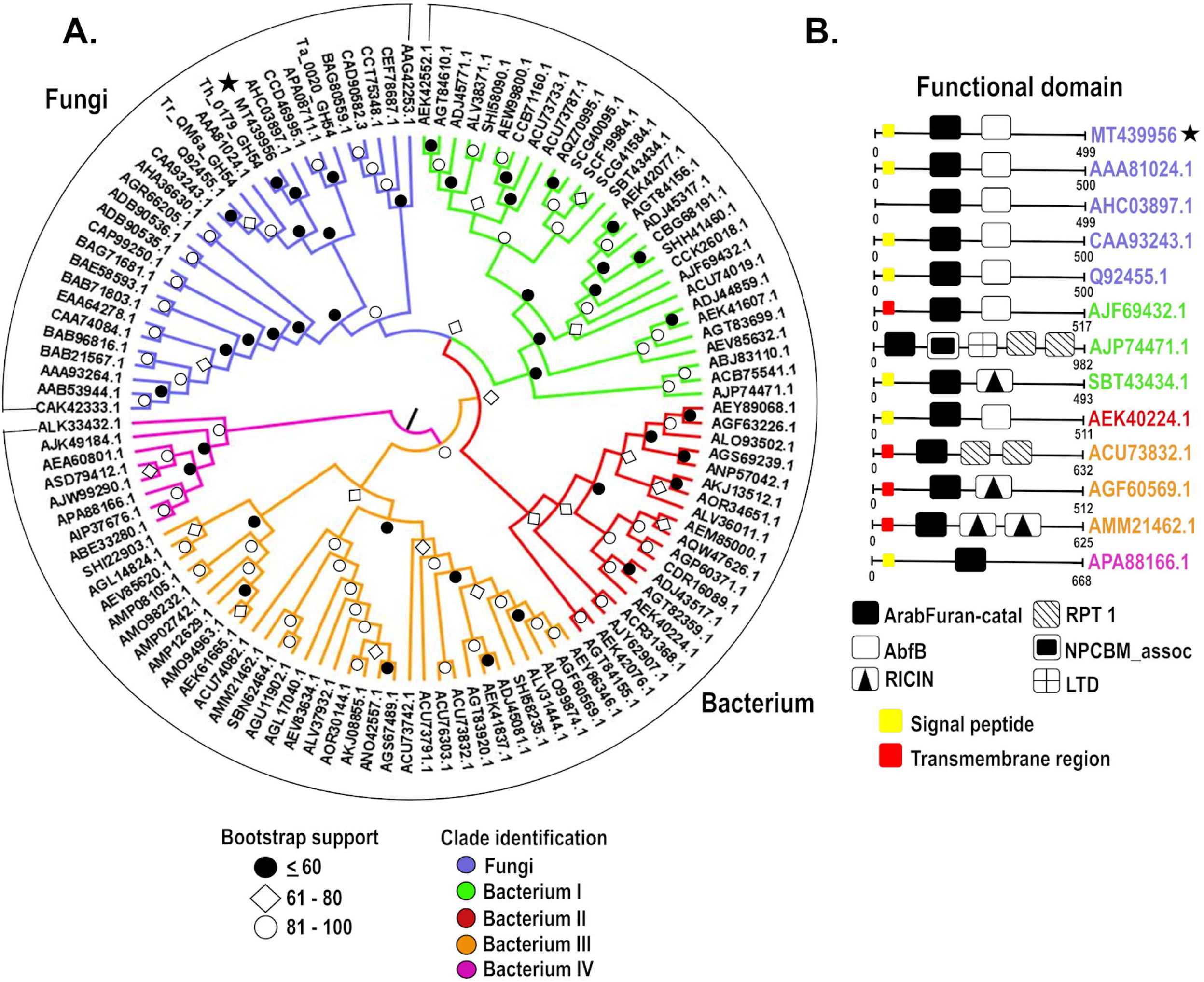
Phylogenetic analysis and sequence domains of the GH54 family. (**a**) Phylogenetic tree of the sequences. The star indicates the studied protein. Below the tree is a legend with the bootstrap values divided into ≤ 60%, 61-80 and 81-100. (**b**) Differences in the composition of domains in the sequences distributed in the groups. The identification of the groups is the same for A and B.

The analysis of the domains of the sequences in the phylogenetic tree (Supplementary Table S2) showed that all members of Fungi contained ArabFuran-catal (α-L-arabinofuranosidase B catalytic) and AbfB (α-L-arabinofuranosidase B) domains, in addition to CBM 42 (Fig. 1b). In Bacterium I, the presence of CBMs 13 and 42 was observed, in addition to sequences with the following combinations of domains: ArabFuran-catal and AbfB; ArabFuran-catal, RICIN (ricin B lectin domain), LTD (lamin tail domain), NPCBM_assoc (NEW3 domain of α-galactosidase) and RPT1; and ArabFuran-catal and RICIN. All Bacterium II sequences exhibited only CBM 42 and ArabFuran-catal and AbfB domains.

The Bacterium III group sequences only CBM 13 and the following combinations of domains: ArabFuran-catal and two RPT1 domains; ArabFuran-catal and RICIN; and ArabFuran-catal and two RICIN domains. Bacterium IV was the smallest group; its sequences included no CBMs and were the most distinct from those of the other groups. The Bacterium IV members included only the ArabFuran-catal domain.

To analyze the similarity of the ThABF sequence with some arabifuranosidases characterized as GH54 members in *Trichoderma*, alignment was performed (Fig. 2a). It was found that the target sequence was most similar to a sequence from *T. virens*, with a percent identity of 92.38%, followed by sequences from *T. reesei* (88.38%) and *T. koniingi* (87.78%). The structure of the ThABF protein was also predicted, and the model that was used was an *Aspergillus kawachii* α-L-arabinofuranosidase B with a C-score of 0.95, a TM-score of 0.84 ± 0.08 and an RMSD of 5.3 ± 3.4 Å (Fig. 2b).

**Figure 2.**
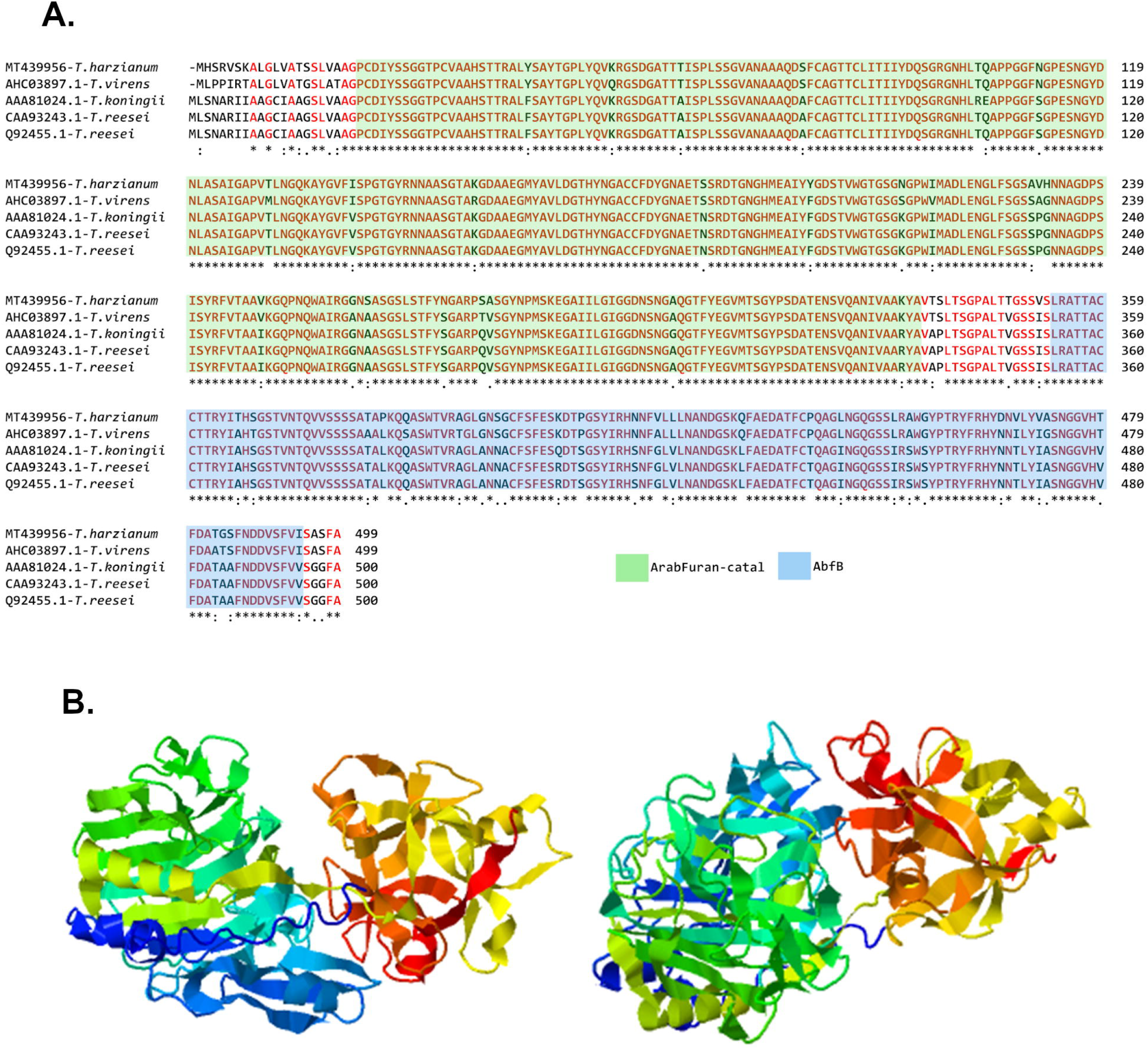
Alignment and structure prediction of ThABF. (**a**) Alignment of ThABF with other *Trichoderma* sequences belonging to the GH54 family. (**b**) Prediction of the ThABF structure by I-TASSER.

### Production of recombinant protein and evaluation of the folding components

With the sequence obtained from ThABF *in silico*, cloning and expression were performed. The enzyme was present in the insoluble fraction (Fig. 3a), and it was necessary to use the refolding approach described in the methodology, through which a protein of 53.44 kDa was purified (Fig. 3b). The recombinant ThABF sequence was used to evaluate the secondary and tertiary folding components of the enzyme since it was obtained through refolding. On the basis of the obtained circular dichroism profile, we identified rich α helices in the ThABF structure (Fig. 3c). In the size exclusion chromatography (SEC), the protein was eluted in a single peak, presenting the characteristics of a monomer in solution (Fig. 3d). Thus, these analyses demonstrated that it was possible to obtain a protein with folding components through the refolding method used.

**Figure 3.**
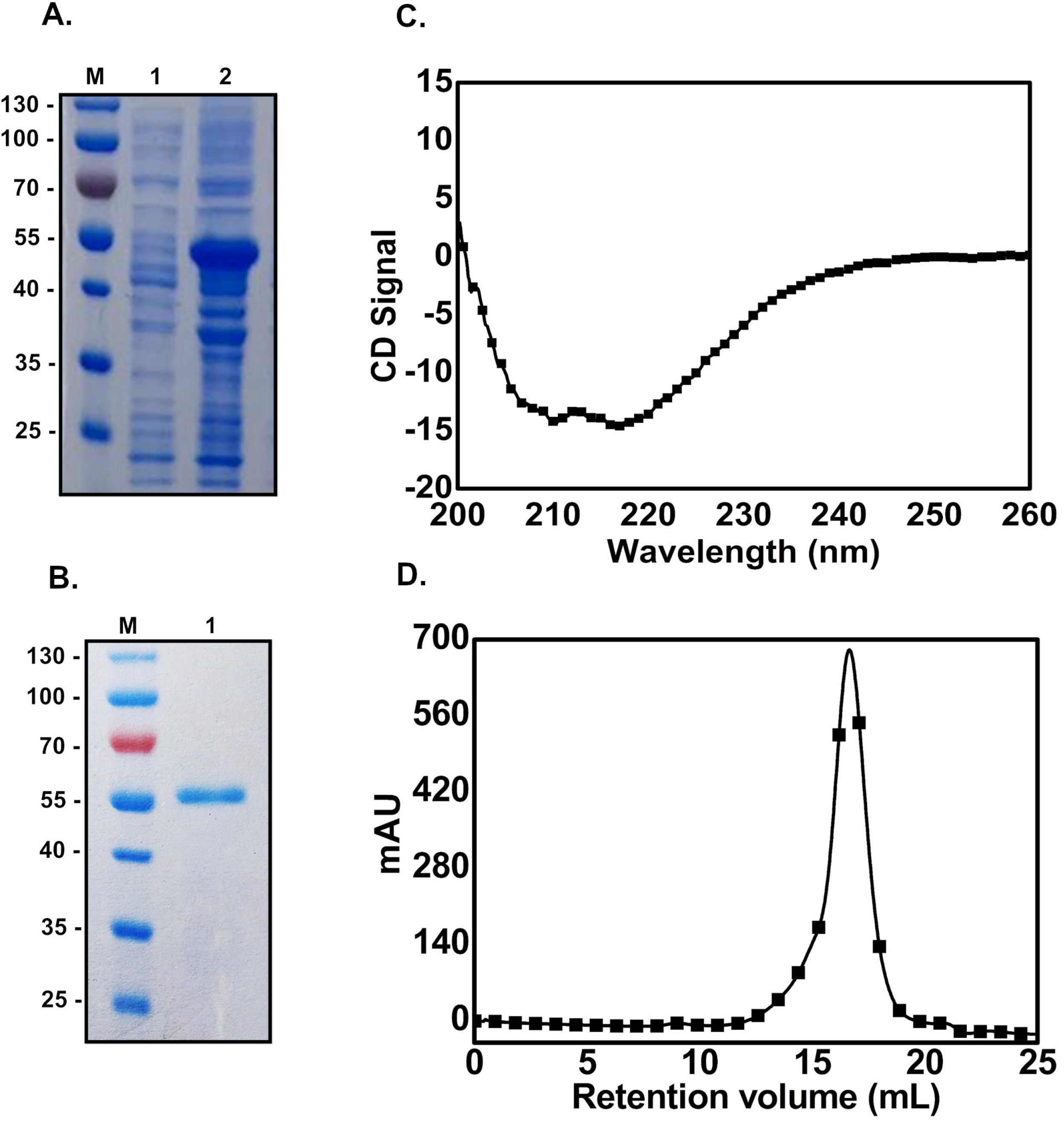
SDS-PAGE in 12% gels showing purification results and graphs of secondary and tertiary folding analyses. (**a**) Gel containing the soluble (1) and insoluble (2) fractions; M indicates the marker. (**b**) Gel containing purified ThABF, with as size of approximately 53.44 kDa (1), and a marker (M). (**c**) Analysis of secondary folding components of the recombinant enzyme by circular dichroism. (**d**) Analytical size exclusion chromatography of ThABF.

### Determination of enzyme specificity and tests with metal ions

In the first experiment, enzyme activity was tested with the pNPAra substrate because *in silico* analysis indicated that ThABF would present a relatively high affinity with this substrate. However, the detected activity was not significant, leading to the hypothesis that a cofactor is needed to activate the catalytic site of the enzyme. Thus, we evaluated the effects of different metals on the activity of ThABF (Fig. 4). It was observed that the enzyme requires a metal ion cofactor to exert its catalytic activity, since the control exhibited a profile similar to that of ethylene diamine tetraacetic acid (EDTA). CUCl2 had no effect on the enzyme’s activity. MgCl2 had the greatest effect on the enzyme’s activity among the eleven tested metal salts, followed by MnCl2, CoCl2, CaCl2 and NiCl2. Despite the demonstration that ThABF requires a cofactor, its activity against α-D-arabinofuranoside (pNPAra)was not significantly increased; thus, a panel of chromogenic substrates derived from nitrophenol was tested to obtain information on ThABF enzymatic specificity (Table 1). The enzyme exhibited the greatest activity against β-D-galactopyranoside (pNPG), followed by α-D-arabinopyranoside (pNPAp), β-D-fucopyranoside (pNPF) and finally pNPAra.

**Table 1.** Relative activity of ThABF against different synthetic substrates.

| Synthetic substrates | Relative activity (%) |
| --- | --- |
| **β-D-galactopyranoside** | $100 \pm 0$ |
| **α-D-arabinopyranoside** | $61.72 \pm 0$ |
| **β-D-fucopyranoside** | $21.79 \pm 0.15$ |
| **α-D-arabinofuranoside** | $4.97 \pm 0.13$ |
| **α-D-glucopyranoside** | $1.12 \pm 0.03$ |
| **α-L-fucopyranoside** | $0.69 \pm 0.60$ |
| **α-L-rhamnopyranoside** | $0.44 \pm 0.53$ |
| **β-D-glucopyranoside** | $0.34 \pm 0.09$ |
| **α-D-xylopyranoside** | $0.26 \pm 0.05$ |
| **α-D-mannopyranoside** | $0.25 \pm 0.05$ |
| **α-D-galactopyranoside** | $0.16 \pm 0.09$ |
| **β-D-cellobioside** | $0.09 \pm 0.04$ |
| **β-D-xylopyranoside** | $0.07 \pm 0.09$ |
| **β-D-mannopyranoside** | $0 \pm 0.05$ |

**Figure 4.**
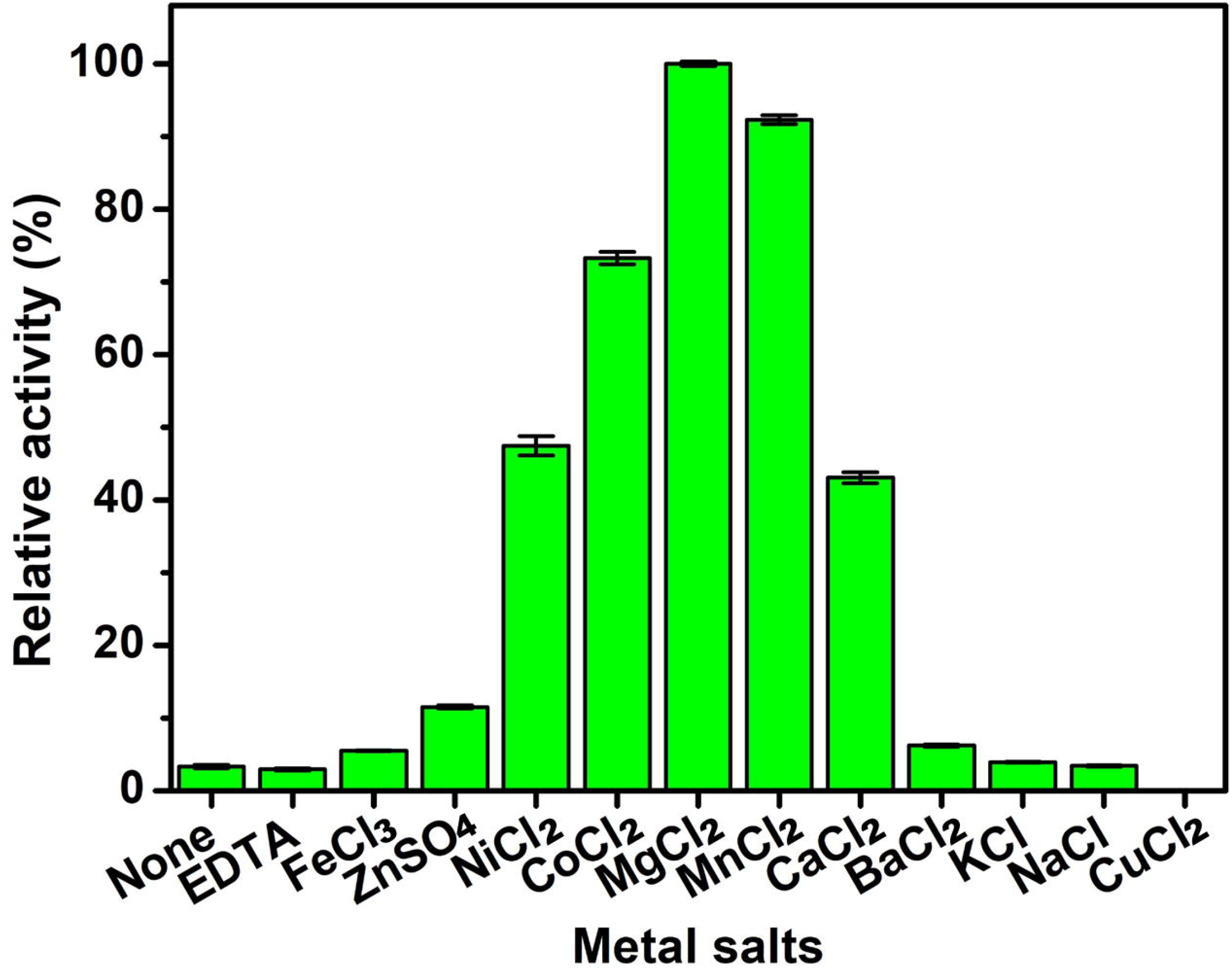
Relative activity of ThABF in the presence of different metallic ions.

### Optimization of pH and temperature conditions and enzymatic kinetics

Following the selection of ThABF-specific substrates and ions, the optimization of pH and temperature was performed under different conditions to identify the ideal performance of the enzyme. As shown in Fig. 5a and b, maximum ThABF activity was observed at 55°C and pH 6.5. The enzyme exhibited greater than 50% activity between temperatures of 45 and 60°C. However, at lower temperatures, in the range of 20 to 40°C, its activity was below 40%. ThABF exhibited a wide range of performance levels at pH values from 6.0 to 9.0, and the optimal pH was shown to be 6.5 in Na2HPO4 phosphate buffer. At pH 4 to 5.5, enzyme activity decreased significantly. Experiments were carried out to examine the kinetics of ThABF with the substrates selected in the specificity tests; the parameters Vmax, Km, Kcat and Kcat/Km were calculated, and the results are shown in Table 2. The curves constructed with the substrates pNPG and pNPAra are shown in Fig. 5c and d.

**Table 2.**
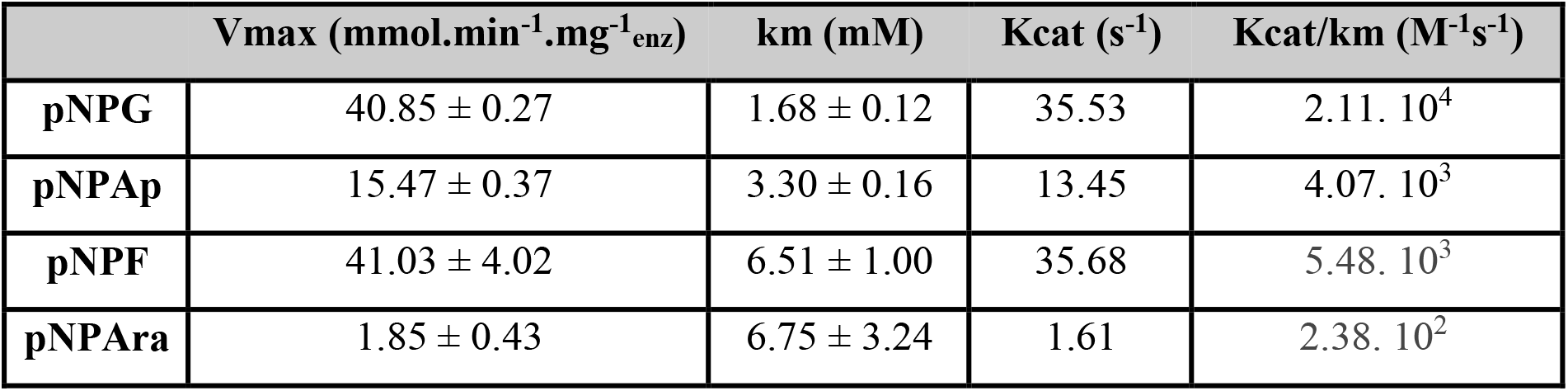
Kinetic parameters of ThABF against selected substrates.

**Figure 5.**
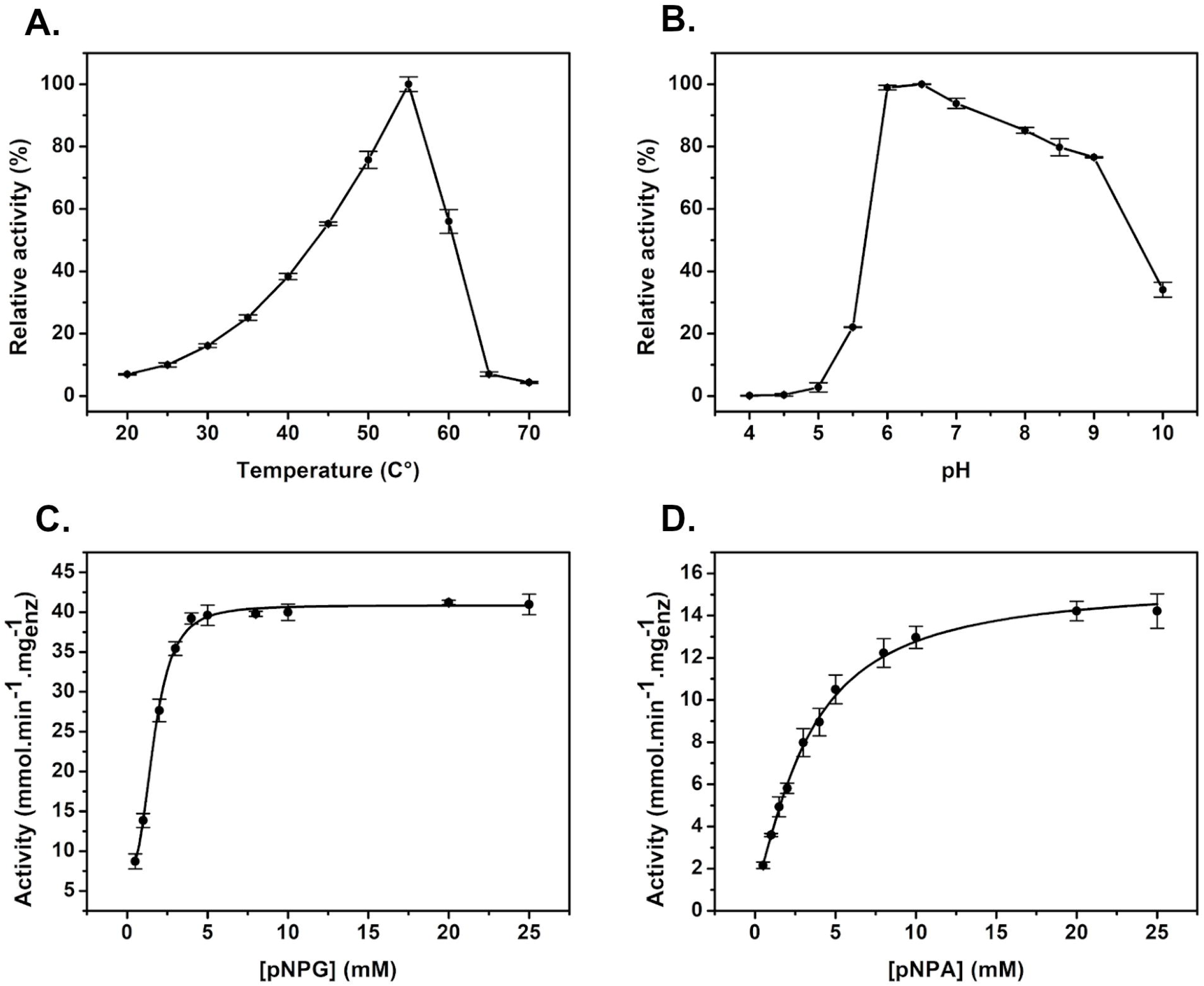
Biochemical characterization of ThABF. Relative activity of the recombinant protein at temperatures of 20 to 70°C (**a**) and in the pH range of 4 to 10 (**b**). Enzymatic kinetics of TiBgal54A with the substrates pNPG (**c**) and pNPA (**d**).

## Discussion

Arabinofuranosidases are enzymes that, in addition to acting on L-arabinose, can also degrade other types of sugars and can be applied in various industrial processes^1,14^. In this work, we bioprospected an arabifuranosidase from the metal-dependent GH54 family from *T. harzianum* and demonstrated that it showed expanded activity against other substrates. This enzyme can be introduced in enzymatic cocktails to assist in the hydrolysis of hemicellulose, since this biopolymer is present in approximately 50% of the most diverse sources of plant biomass, and its degradation requires highly diverse enzymes due to its structural complexity^15–18^.

Through the *in silico* analyses carried out in this study, great functional diversity was revealed in the GH54 family. Some studies have demonstrated that enzymes of this family exert activity against other types of substrates, in addition to those that are commonly identified, such as β-D-galactofuranoside, α-L-arabinopyranoside and β-D-xylopyranoside^4,19^. Evidence supporting this conclusion was found in a phylogenetic analysis in which we observed the presence of LTD, RICIN and NPCBM domains (related to the cleavage of simple sugars such as galactose and lactose) in several not yet characterized sequences^20–23^. We also identified several types of CBMs related mainly to enzymes that act on hemicellulose from both the phylogenetic and BLAST results. CBMs 42 and 13 are the most prevalent in GH54 sequences, possibly because they are present in some arabinofuranosidases and xylanases^24,25^.

Through the alignment and prediction of the structure of ThABF, we showed that the target protein identified in this study has the characteristics of an arabinofuranosidase, particularly regarding the high conservation of amino acids in the ArabFuran-catal and AbfB catalytic domains^24,26,27^. The ArabFuran-catal domain was present in all sequences analyzed in the family. This domain targets several L-arabinose bonds present in hemicellulose, as does AbfB^24,26^. For the purification of ThABF, it was necessary to use a refolding method adapted to recover the protein from inclusion bodies in its best oligomeric state^28^. The final protein yield following purification was relatively low, at 0.51 mg/ml, which made it difficult to carry out other types of biochemical tests; however, the method allowed us to characterize the enzyme and study it, as demonstrated in other studies^28,29^.

In the tests with different substrates, we demonstrated that ThABF exerted activity on the substrates pNPG, pNPAp, pNPF and pNPAra. As previously described, some GH54 enzymes exhibit activity against substrates other than those reported in this family; however, there have been only two studies addressing this topic^4,19^. At the time of writing the present report, 19 enzymes had been characterized in this family, among which only 6 have been tested on different types of synthetic substrates^4,19,30,31^. In addition, as demonstrated via *in silico* analyses, there are several unstudied proteins of this family harboring domains related to the hydrolysis of galactose, among other types of sugars. This demonstrates the limitation of studies involving enzymes of this family of GHs.

The data obtained in the tests with metallic cofactors showed that the enzyme was dependent on metals. This role of Mg^2+^ has been reported for other enzymes; there are several hypotheses regarding how magnesium acts on these enzymes, one of which is that it helps maintain the conformation of the active site^32^. The analysis of the optimal conditions of the enzyme showed that ThABF presents characteristics similar to those of other fungal arabifuranosidases of the GH54 family, making it quite adaptable to changes in the conditions imposed in several types of biotechnological processes, such as the hydrolysis of plant biomass^1^.

Based on the results presented above, ThABF has the potential to be used in the saccharification of lignocellulosic material, mainly because it acts on different types of sugars that constitute the side chains of hemicellulose^33,34^. These side chains contribute to the recalcitrancy of plant biomass due to its diverse structural composition^2,35^. The removal of hemicellulose makes it less recalcitrant, improving the hydrolysis process^8,9^. For this purpose, enzymes such as arabinofuranosidases that act on these sugars are necessary^34^.

In conclusion, the present study reports the *in silico* bioprospection, expression, purification and biochemical characterization of a GH54-dependent metal with expanded substrate specificity that acts in a wide pH range. These characteristics indicate the potential to use this enzyme in industrial bioprocesses, such as the saccharification of plant biomass.

## Methods

### In silico *data mining, RNA-Seq read mapping and phylogenetic analyses of ThABF*

A search for sequences of bifunctional *Trichoderma* enzymes that act on hemicellulose was carried out in the NCBI nonredundant protein^36^, UniProt and CAZy databases. With the selected sequence, a BLASTP search was performed versus the genome of *T. harzianum* CBS 226.95 (TaxID: 983964) in the NCBI database. The most similar sequence was used to map the RNA-Seq reads of the *T. harzianum* IOC-3844 data generated in the work of Horta *et al*.^37^ and, thus, obtain the sequence of the specific protein of the target lineage. For mapping, CLC *Genomics Workbench software* (CLC bio - v4.0; Finlandsgade, Dk) was used. Analyses of physical-chemical parameters were also performed with ProtParam; the presence of signal peptides was evaluated on the SignalP-5.0 server; domain prediction was conducted with SMART; and protein structure prediction was conducted with I-TASSER.

A BLASTP search of the sequence resulting from mapping was performed versus the CAZy database, and different enzymes of the GH54 family were then selected to perform phylogenetic analysis. The sequences were aligned using *ClustalW*^38^ implemented in *Molecular Evolutionary Genetics Analysis software*, version 7.0^39^. Gaps were removed manually, and incomplete or difficult-to-align sequences were excluded from the analysis. Phylogenetic analyses were performed with *MEGA7* using *maximum likelihood* (ML)^40^ inference based on the Jones-Taylor-Thornton (JTT) model with 1000 *bootstrap*^41^ repetitions for each analysis. The tree, drawn to scale, was obtained automatically by applying the *neighbor-joining* BioNJ algorithms to a matrix of distances in pairs estimated using the JTT model and the topology with the highest log probability values. The trees were visualized and edited using the program *Figtree*^42^.

### Heterologous production and refolding of TiBgal54A

The ThABF gene was cloned into the pET-28a (+)vector using the standard gene cloning protocol of Sambrook^43^ with the aid of the forward primer TAAGAATTCGGGCCCTGTGA and the reverse primer TGGTCGACTTAAGCAAACTGG. The recombinant plasmids were transformed into *Escherichia coli* Rosetta (Novagen, Darmstadt, Germany) for protein overexpression. Transformed *E. coli* Rosetta cells were grown to an optical density of approximately 0.8 at 800 nm. To trigger the transcription of the ThABF gene, it was inserted in a culture containing isopropyl β-D-1-thiogalactopyranoside (IPTG) at 0.4 mM. Then, the cells were incubated overnight at 25°C, harvested by centrifugation and resuspended in 50 mL of buffer A (250 mM NaCl, 40 mM sodium phosphate, 20 mM imidazole, 3 mM MgCl2, 1 mM EDTA and pH 8.0), 1 mg/mL lysozyme and 1 mM phenylmethylsulfonyl fluoride (PMSF) for 30 min with shaking on ice. The cells were disrupted by sonication, and the soluble fraction was obtained by centrifugation (16,000 rpm, 40 min, 4°C).

After rupturing the cells and resuspending the inclusion bodies, the refolding protocol of Santos *et al*.^28^ was applied. The pellet was resuspended in 15 mL of buffer A containing 1 M urea. Then, sonication and centrifugation (16,000 rpm, 15 min, 4°C) were performed 6 times. Dialysis was subsequently performed under agitation *overnight* with buffer A plus 0.4 M arginine, followed by centrifugation (10,000 rpm, 10 min, 4°C). The concentrations of the purified proteins were determined spectroscopically using the molar extinction coefficient (ε) predicted on the basis of the amino acid sequence, and sample purity was estimated by polyacrylamide gel electrophoresis with 12% sodium dodecyl sulfate (SDS-PAGE).

### Analysis of folding components

The far-UV CD spectra of ThABF were collected using a Jasco model J-810 spectropolarimeter from the National Biorenewables Laboratory (LNBR) coupled to a Peltier control system (PFD 425S-Jasco). The CD spectra were generated using the purified recombinant protein at a concentration of approximately 1.024 mg/ml in 10 mM sodium phosphate buffer, pH 8.0. A total of 12 accumulations were recorded within the range of 260 to 208 nm at a rate of 50 nm min^−1^ using a quartz cuvette with a travel length of 1 mm, and the results were averaged.

The tertiary folding components of the recombinant refolded ThABF were evaluated by analytical SEC using a Superdex 200 10/300 GL column (GE Healthcare, Uppsala, Sweden). The protein sample at a concentration of approximately 0.56 mg/ml was dialyzed against buffer A, and gel filtration was then performed at a flow rate of 0.5 mL.min^−1^. The elution fractions from each chromatographic run were collected and analyzed by 12% SDS-PAGE.

### Biochemical characterization

The substrate specificity of ThABF was tested with the following pNP glycosides: α-D-arabinopyranoside (pNPAp), β-D-galactopyranoside (pNPG), α-D-arabinofuranoside (pNPAra), β-D-fucopyranoside (pNPF), α-D-glucopyranoside, α-D-xylopyranoside, α-D-mannopynoside, α-L-ramnopyranoside, α-L-fucopyranoside, β-D-glucopyranoside, α-D-galactopyranoside, β-D-xylopyranoside, β-D-cellobioside and β-D-mannopyranoside. A 10 mM concentration of pNPs was used in the tests, which were conducted in 100 mM phosphate buffer (HPO_4_), pH 6.5, at 55°C for 10 min. The reactions were stopped with 100 mM calcium carbonate (CaCO_3_), and the released pNP was quantified spectrophotometrically at 410 nm.

After selecting the substrates against which ThABF exhibited activity, its activity was tested with different 1 mM concentrations of different metal salts (CaCl_2_, KCl, NaCl, MgCl_2_, MnCl_2_, CoCl_2_, ZnSO_4_, CuCl_2_, NiCl_2_, FeCl_3_ and BaCl_2_) and EDTA. Before these tests, any metal ions that could interfere with the analysis were removed via two dialysis steps performed on the recombinant protein overnight under agitation. First, 15 mM EDTA was added to chelate the metals present in the sample, and the protein was then transferred to a buffer containing 10 mM sodium phosphate. Subsequently, the tested chemicals were individually incubated with the recombinant enzyme; the reactions occurred at 55°C at pH 6.5 for 10 min with pNPG as the substrate.

After the selection of the ion with the greatest effect on the enzyme, it was used to optimize pH and temperature conditions. The optimal pH was determined by incubating 0.12 mg/ml of purified ThABF with 10 mM pNPG in buffers with pH levels ranging from 3.0 to 10.0 [CH3COONa sodium acetate buffer (100 mM, pH 3.0 to 5.0), Na2HPO4 phosphate buffer (100 mM, pH 6.0 to 8.0) and Tris-NaCl (100 mM, pH 7.0 to 9.0)]. The mixtures were incubated at 55°C for 10 min, and the released pNP was quantified spectrophotometrically at 410 nm. The same conditions were applied to investigate the ideal temperature; these tests were conducted with Na2HPO4 phosphate buffer (pH 6.5, 100 mM) at a temperature range of 20 to 70°C. Subsequently, the pH test was repeated at the identified temperature.

After performing the above tests, assays of enzymatic kinetics were performed at pH 6.5 at 55°C for 10 min, with pNPG, pNPAp, pNPF and pNPAra as substrates. The parameters Km, Vmax, and Kcat and the Kcat/Km ratio were obtained by plotting using the Lineweaver-Burk method, where the plots were constructed by plotting the substrate concentration on the x axis and the speed of the enzymatic reaction on the y axis using the OriginPro 8.5.0 program. The experiments were performed in triplicate at an ideal temperature and pH. A unit of enzymatic activity was defined as the amount of enzyme required to release 1 μmol of p-nitrophenol per minute under the tested conditions.

## Acknowledgements

This work was supported by grants from the Fundação de Amparo à Pesquisa do Estado de São Paulo (FAPESP 2015/09202-0 and 2018/19660-4), Coordenação de Aperfeiçoamento de Pessoal de Nível Superior (CAPES) Computational Biology Program (CBP: Process number 88882.160095/2013-01) and Conselho Nacional de Desenvolvimento Científico e Tecnológico (CNPq: Process number 312777/2018-3). MLLM received a Ms fellowship from CAPES (CBP - 88887.200427/2018-00); JAFF received a PhD fellowship from CNPq (170565/2017-3) and a PD fellowship from CAPES (CBP - 88887.334235/2019-00); CAS received a PD fellowship from FAPESP (2016/19775–0) and an SWE PD fellowship from CAPES (CBP); LMZ recipient of a research fellowship from FAPESP (19/08855-1); RRM received a PD fellowship from FAPESP (17/14253-9); and APS is the recipient of a research fellowship from CNPq (312777/2018–3). We thank the National Biorenwables Laboratory (LNBR).

## Author contributions

M.L.L.M. carried out all the experiments and wrote the manuscript. J.A.F.F. helped with the *in silico* analyses and wrote the manuscript. R.R.M. and L.M.Z. helped with the biochemical characterization. C.A.S. designed the experiments and wrote the manuscript. A.P.S. directed the general study and wrote the manuscript. All authors read and approved the manuscript.

## Accession Codes

The protein sequence studied in the study is deposited at the national center for biotechnology information (NCBI) with the access code MT439956.

## Additional information

**Supplementary Table 1.** ThABF Blastp against CAZy bank.

**Supplementary Table 2.** GH54 sequences used in the phylogenetic tree.

